# Epigenetic regulation of TERRA transcription, R-loop formation, telomere integrity and metacyclogenesis by base J in *Leishmania major*

**DOI:** 10.1101/2024.06.27.601056

**Authors:** Rudo Kieft, Robert Sabatini

## Abstract

The hyper-modified DNA base J helps control termination of Pol II transcription at polycistronic transcription units (PTUs) in *T. brucei* and *L. major*, allowing epigenetic control of gene expression. The Telomere Repeat-containing RNA (TERRA) is synthesized in *T. brucei* by Pol I readthrough transcription of a telomeric PTU. While little is understood regarding TERRA synthesis and function, the hyper-modified DNA base J is highly enriched at telomeres in *L. major* promastigotes. We now show that TERRA is synthesized by Pol II in *L. major* and loss of base J leads to increased TERRA from multiple telomeric ends, presumably via readthrough transcription from an adjacent PTU. Furthermore, Pol II readthrough defects and increased TERRA correlate with increased telomeric DNA damage, increased differentiation of promastigotes to the infectious metacyclic life stage and decreased cell viability. These results help explain the essential nature of base J in *Leishmania* and provide insight regarding epigenetic control of RNA expression, telomere stability and parasite development during the life cycle of *L. major*.

## INTRODUCTION

Trypanosomatids are a group of early-diverged eukaryotes responsible for multiple diseases including African sleeping sickness and leishmaniasis. The protozoan parasites, which include *Trypanosoma brucei* and *Leishmania major*, progress through life stages by cycling between an insect vector and a mammalian host. Thus, gene expression control is needed for the parasites to rapidly adapt to fluctuating conditions in their environment. Unlike most eukaryotes, the genome of trypanosomatids is composed of genes ordered into long polycistronic transcription units (PTUs) that are transcribed from a single promoter at the 5’ end of the cluster (1–4). The arrangement of functionally unrelated genes into co-transcribed gene clusters has made it difficult to envision regulated gene expression at the level of RNA polymerase II (Pol II) transcription in trypanosomatids. However, recent work characterizing the mechanism and epigenetic control of Pol II transcription termination in *T. brucei* and *L. major* demonstrated a mechanism for control of individual gene expression within a polycistronic system (5–9).

Termination of Pol II transcription in humans and yeast is by the PTW/PP1 complex that includes protein phosphatase 1 (PP1), PP1 interactive-regulatory protein (PNUTS) and the DNA binding protein Tox4 (10–17). Critical to this process is PP1-dependent dephosphorylation of Pol II leading to decreased Pol II elongation, enhancing the capture by the torpedo RNA exonuclease allowing Pol II dissociation and termination (10,12,17,18). A similar PJW/PP1 complex was identified in *T. brucei* and *L. major* and shown to be involved in Pol II termination at the end of PTUs genome-wide via PP1-PNUTS mediated dephosphorylation of Pol II (5,7,19,20). PP1-PNUTS is presumably recruited to termination sites via association with the base J-DNA binding protein, JBP3 (5,19). Base J consists of a glucosylated thymidine (21) found in the nuclear DNA of trypanosomatids at Pol II transcription termination sites (22,23) and shown to represent an epigenetic mark involved in Pol II transcription termination in *T. brucei* and *L. major* (8,9,23–26). J biosynthesis occurs through the hydroxylation of thymidine in DNA by the thymidine hydroxylase JBP1 or JBP2 (27), forming hydroxymethyluridine, followed by the transfer of a glucose by the glucosyltransferase enzyme JGT (28–31). Addition of dimethyloxalylglycine (DMOG) to the growth medium inhibits hydroxylase activity of JBP1/2 and thus, decreases base J levels in the parasite genome (27). While base J is not essential in *T. brucei* (22,32), it appears to be essential in *Leishmania* since JBP1 KO cells are not viable (33) and reduction of base J using DMOG can lead to a severe growth phenotype (9). Reduction of J in *Leishmania* and *T. brucei* using genetic KO of JBPs or via DMOG led to readthrough transcription at termination sites genome-wide, identical to termination defects seen in PP1-PNUTS complex mutants linking base J with dephosphorylation of Pol II, suggesting a critical role for J in transcription termination (8,23–26). Histone variant H3V co-localizes with J at termination sites in *T. brucei* (22) and *L. major* (9) and both epigenetic marks function to promote termination, such that the combined loss of J and H3V results in a synergistic increase in read through transcription (8,9,25,26). For several PTUs in *T. brucei* and *L. major*, J/H3V are found to promote Pol II termination prior to the end of the gene cluster, leading to silencing of the downstream genes (8,9,24). Loss of J/H3V, or components of the PP1-PNUTS complex results in readthrough transcription and de-repression of the downstream genes (5–9,19). This has suggested ‘premature’ termination of Pol II transcription at the end of a trypanosomatid PTU as a novel mechanism of regulating the transcription of specific genes. Genes regulated by Pol II termination include genes that are developmentally regulated and code for proteins involved in optimal growth during infection of the mammalian host (9,26). Therefore, de-repression of specific protein coding genes by Pol II readthrough in *L. major*, and associated *in vitro* growth defects, is thought to explain the essential nature of J in Leishmania (9).

Base J synthesis is developmentally regulated in *T. brucei*, *T, cruzi* and *Leishmania* (34–37). However, *T. brucei* is unique where J is restricted to a single life stage; the infective mammalian bloodstream form where approximately 50% of base J localizes to telomeres (36,38). The most telomere-proximal PTU in some chromosomes of *T. brucei* represent the bloodstream form VSG expression site (VSG ES) involved in regulated expression of the trypanosome surface antigen gene, VSG. Allelic exclusion of VSG expression and periodic switching of the expressed allele allows trypanosomes to evade the host immune system, in a process known as antigenic variation. The VSG ESs are transcribed by Pol I. Only one Pol I promoter is transcriptionally active at a time, while the others are repressed, allowing monoallelic VSG expression. J is found in the silent VSG ESs and is highly enriched in the telomeric repeats, replacing ∼14% of T in (CCCTAA)_n_ and ∼36% in (TTAGGG)_n_, and repeats upstream the Pol I promoter (35,36). The Pol II readthrough defects seen in base J/H3V mutants (8,25,26) and PP1-PNUTS complex mutants (5,7) are associated with derepression of the silent VSG ES. Rather than an independent function of base J controlling Pol I transcription of the telomeric VSG ESs, the TbPP1 KD indicates Pol II termination defects at the chromosome core leads to Pol II extending into the Pol I transcribed telomeric PTUs (7). These results suggest a key function of base J in BS form *T. brucei* is to control pervasive Pol II transcription and help maintain monoallelic expression of VSG.

Monoallelic control of VSG expression in *T. brucei* extends to the downstream telomeric repeat that is transcribed into telomere repeat-containing RNA (TERRA), which is prone to form RNA:DNA hybrids (R-loops) with the telomeric DNA (telomere R-loops or TRLs)(39). Studies in mammalian cells and yeast indicate TERRA (and TRLs) as a key player of telomere biology and genome integrity (39,40). Formation of R-loops can lead to DNA damage caused by induced double-strand DNA breaks (DSBs) and recombination, threatening genome integrity (41–43). In *T. brucei*, TERRA has been shown to form TRLs (44) and higher levels of TERRA and TRLs were associated with higher levels of telomeric and subtelomeric dsDNA breaks and enhanced rate of recombination based VSG switching (44–46). TERRA is transcribed in bloodstream form *T. brucei* by Pol I as a readthrough product, or transcription run-off, when the enzyme continues into the telomeric repeats downstream of the VSG in the active ES I (7,44,47–49). Transcription of TERRA by Pol I is unique to trypanosomes. TERRA is transcribed by Pol II in mammals and yeast (50,51) and subtelomeric promoters identified (52–54). Therefore, in contrast to human and yeast cells where multiple telomeres can be transcribed by Pol II (55), TERRA is transcribed in *T. brucei* from a single telomere and as a transcription run-off product by Pol I. However, transcription of TERRA from the active ES is increased by Pol II readthrough transcription in the TbPP1 mutant (7). These results indicate that the control of Pol II termination, involving base J, helps control the levels of TERRA and telomere biology in BS form *T. brucei*.

In the *L. major* life cycle the parasites transform into three main morphologically distinct developmental stages (amastigote, promastigote, and metacyclic promastigote). Amastigotes that reside in mammalian macrophages transform into promastigotes in the alimentary tracts of their invertebrate vector (sand flies). Promastigotes multiply in the insect midgut, migrate to the insect’s stomodeal valve and differentiate into the infective non-dividing metacyclic promastigote form (56). This differentiation process, called metacyclogenesis can be replicated in vitro in axenic culture (57). Here, while log phase procyclic promastigotes enter the stationary phase a fraction of the parasites differentiate into metacyclics. While a fraction of base J is localized and functional at Pol II termination sites within the *Leishmania* genome, DNA hybridizations indicate >90% of the total J is localized at telomeres (58), suggesting an important telomere function. Analysis of blocked restriction digestion suggested developmental regulation of telomeric J in *L. major* with reduced levels in metacyclics (37). While little is known regarding TERRA synthesis and function in Leishmania, it has been shown to be developmentally regulated with increased levels in infective metacyclics and potentially synthesized from multiple telomeric ends (37,48). While no Pol II promoter has been described, RT-PCR analysis of specific telomeric ends in *L. major* have detected TERRA synthesis in chromosomes with a PTU transcribed towards the telomere as well as those that have a unit directed away from telomeres (48). This has suggested TERRA is the product of readthrough (or run-off) Pol II transcription of telomeric PTUs as well as subtelomeric localized promoters. However, the lack of epigenetic markers that represent transcription start sites in *Leishmania* does not support TERRA specific subtelomeric promoters and PRO-seq analysis of transcription in *L. major* supports Pol II run-off transcription at ends with a PTU transcribed towards the telomere (59). Base J regulation of Pol II termination and parasite growth in *L. major* promastigotes, along with the apparent modulation of telomeric J and TERRA transcript levels throughout the life cycle, suggest a J regulatory mechanism of TERRA expression and an essential stage-specific role in these early-branching eukaryotes.

Here we set out to determine the function of base J in regulating TERRA expression in *L. major*. Our data indicate that TERRA is synthesized by Pol II in *L. major*, primarily in chromosomes with a PTU transcribed towards the telomere, and loss of base J leads to increased TERRA, independent of detectable change in TERRA stability. Modulating base J levels in vivo using DMOG inhibition of thymidine hydroxylase activity and deleting H3V can lead up to ∼100-fold increase in TERRA. RT-PCR analysis indicates this is primary due to Pol II readthrough/run-off transcription into telomeric repeats from the adjacent PTU at multiple chromosome ends. Depletion of J also led to increased levels of R-loops and DSBs at telomeres. Therefore, base J control of Pol II transcription termination regulates TERRA synthesis and TRL levels, the latter of which controls telomeric DNA stability. The increased TERRA in base J mutants is also associated with an increase in metacyclogenesis and decreased cell viability. The phenotypes we describe here suggest that epigenetic control of Pol II termination and TERRA synthesis may explain the essential nature of base J in Leishmania. The impact of these studies on understanding to the biological role of the unusual non-coding TERRA RNA in *Leishmania* will be discussed.

## MATERIALS AND METHODS

### Parasite culture

Promastigote *Leishmania major* strains, wild type and the H3V KO, were grown in M199 medium, supplemented with 10 % FCS at 26°C as previously described (9). DMOG treatment of cells was performed by supplementing media with 2.5 mM or 5 mM DMOG for 5 days. Control cells were treated with an equal amount of vehicle (DMSO).

### Metacyclogenesis

We used the lectin peanut agglutinin (PNA) protocol to purify metacyclic parasites from stationary-phase promastigote of *L. major* (60–62). *L. major* promastigotes are agglutinated by peanut lectin while metacyclics are not. Briefly, promastigotes were grown for 7 days to reach stationary phase. Parasites were then harvested at 2000g for 10 min, washed and resuspended in complete medium at 2 × 10^8^ cells/ml. Peanut lectin was added to a final concentration of 50ug/ml and incubated at room temperature for 30 min. Agglutinated parasites (promastigotes) were cleared by centrifugation at 200g for 10 min. Metacyclic stage parasites (PNA-) were recovered from the supernatant by centrifugation at 2000g, resuspended in PBS and counted.

### Determination of the genomic level of base J

To quantify the genomic level of base J, we used the anti-J DNA immunoblot assay as described (32,36) on total genomic DNA, which was isolated as described (63). Briefly, equal amounts of serially diluted genomic DNA was blotted to nitrocellulose followed by incubation with anti-J antisera. Bound antibodies were detected by a secondary goat anti-rabbit antibody.

For quantitating the levels of base J in telomeres, genomic DNA was sonicated and anti J immunoprecipitation was performed as previously described (22). Each J-IP experiment was performed in triplicate and immunoprecipitated J containing DNA analyzed by telomere quantitative PCR. Input DNA was used as a positive control for PCR.

### Nuclear run-on

1.5 × 10^8^ cells were washed 2x in ice cold PBS and washed 1x in ice cold lysis buffer (10 mM Tris pH 7.5, 3 mM CaCl_2_, 2 mM MgCl_2_) and resuspended in 10 ml lysis buffer. NP40 was added to a final concentration of 0.5%, transferred to a Dounce homogenizer and broken with 50 strokes with a tight pestle. Nuclei were harvested after centrifugation for 10 min at 1400 x g, washed with 10 ml lysis buffer and resuspended in 50 microliter lysis buffer. Nuclei were then incubated after addition of H_2_O or α-amanitin at a final concentration of 1 mg/ml for 10 min on ice and 5 min at 30°C. Nuclei were pelleted and resuspended in 100 microliter transcription mixture (100 mM Tris pH 7.5, 25 % glycerol, 0.15 mM Spermidine, 0.5 mM Spermine, 2 mM DTT, 2 mM MgCl_2_, 4 mM MnCl_2_, 50 mM NaCl, 50 mM KCl, 2 mM ATP, 2 mM GTP, 2 mM CTP, 10 μM UTP, 2 mM 5-Ethynyl Uridine, 50 U Superase-In and 10 μg / ml Leupeptin) in the presence or absence of 1 mg/ml α-amanitin. After incubation for 20 min at 30°C nuclei were lysed in TriPure Isolation Reagent. RNA was prepared and 3 μg was subjected to click chemistry (ThermoFisher, C10365) according to the manufacturer’s instructions. Briefly, incorporated EU residues were biotinylated and purified with magnetic streptavidin beads. cDNA was generated with bound biotinylated RNA on the beads as described below with strand specific oligonucleotides. WT cells have the blasticidin gene inserted into the rRNA locus (6), providing a Pol I transcribed control.

### Northern analysis

Total RNA was isolated with Tripure Isolation Reagent. For Northern blot analysis, 10 μg of Turbo DNase treated total RNA was separated on a 1.2 % agarose gel (1 x MOPS, 2.8 % formaldehyde), transferred to a nitro cellulose filter in 20 X SSC and UV cross-linked. Radiolabeled oligonucleotides were generated and hybridized in an aqueous hybridization buffer with 100 μg/ml yeast tRNA at 60°C. A random primer labeled β-Tubulin probe was generated and hybridized in a 40 % (v/v) formamide hybridization mix with the addition of 10% (w/v) dextran sulfate and 100 μg/ml salmon sperm DNA at 42°C. Final washes were performed with 0.3 X SSC / 0.1 % SDS at 60°C for 30 min.

### Quantitative RT-PCR analysis

Total RNA was isolated with Tripure Isolation Reagent (Roche). cDNA was generated from 0.5-1 μg Turbo^TM^ DNase (ThermoFisher) treated total RNA with Superscript^TM^ III (ThermoFisher) according to the manufacturer’s instructions with either oligo dT primers or strand specific oligonucleotides. Quantification of Superscript^TM^ III generated cDNA was performed using an iCycler with an iQ5 multicolor real-time PCR detection system (Bio-Rad). Triplicate cDNA’s were analyzed and normalized to tubulin cDNA. qPCR oligonucleotide primers combos were designed using Integrated DNA Technologies software. cDNA reactions were diluted 20-fold and 5 µl was analyzed. A 15 µl reaction mixture contained 4.5 pmol sense and antisense primer, 7.5 µl 2X iQ SYBR green super mix (Bio-Rad Laboratories). Standard curves were prepared for each gene using 5-fold dilutions of a known quantity (100 ng/µl) of WT gDNA. The quantities were calculated using iQ5 optical detection system software.

Quantification of telomeric DNA in a sample was performed using a telomeric primer pair that specifically amplifies telomeric hexamer repeats without generating primer dimer-derived products (64). Additional bases at the 5’ end of each primer also blocks them from initiating DNA synthesis in the middle of the PCR product in subsequent cycles.

### TERRA RT-PCR analysis

For the semi-quantitative RT-PCR analysis to determine the origin of TERRA, total RNA was isolated and Turbo^TM^ DNase treated as described above, except DNase I digestions were repeated two additional times. Reverse transcription was performed using TELC20 as the primer and PCR amplified. To reduce endogenous priming by the telomeric primer TELC20 in the RT reaction, the denaturation step was performed by heating to 90°C for 1 min followed by cooling to 55°C at 0.5 °C / sec. For PCR, equal amounts of (strand specific) cDNA was used with Ready Go Taq Polymerase (Promega).

qPCR data for TERRA quantification were analyzed using the relative quantification method. Dissociation curves were included in all amplification runs, and Ct (threshold cycle) values were calculated. The tubulin gene was used as a reference gene to normalize the gene expression data. The relative level of TERRA expression was calculated using the 2^−ΔCt^ method (65–69), obtained through the difference in threshold cycle between TERRA and the tubulin reference gene. Tubulin specific primer was utilized in the RT step. Analyses were performed by calculating the averages and SDs of the 2^−ΔCt^ values, normalized for tubulin mRNA, for each triplicate.

### DNA-RNA hybrid immunoprecipitation and qPCR (DRIP-PCR)

Approximately 2×10^7^ cells of *L. major* log-phase promastigotes were used to obtain cross-linked sonicated chromatin as previously described (9). The final nucleic acid suspension was divided into two aliquots. One aliquot was left untreated, and the other was treated with 20 U of recombinant *Escherichia coli* RNase HI (Invitrogen and Thermo Fisher Scientific) overnight at 37°C. Nucleic acids were extracted with an equal volume of ultrapure phenol:chloroform:isoamylic alcohol (25:24:1) and precipitated with 0.1 V 3M sodium acetate pH5.2 and 2 V of ice-cold 100% ethanol. Pellets were suspended in 0.1 mL of ultrapure TE pH8.0 and 5% of the final solution was saved as input. 40 ug of DNA was then immunoprecipitated with the S9.6 antibody or mouse IgG as a control as previously described (70).

DNA obtained from immunoprecipitated chromatin was subjected to qPCR reactions as described above, using primers to assay telomeric DNA and specific internal regions of the genome on chromosome 36 and 7 previously demonstrated to contain differing levels of R-loops (70). The LmjF.07.0250 ORF (7.025) is a R-loop negative region and UTR regions in chromosome 36 (Ch36A and Ch36B) were shown to be R-loop positive. ChIP enrichment of R-loops at each region were calculated using Ct values, using the formula 100×2^(adjusted input – Ct (IP))^ with adjusted input = Ct (input) – log_2_(20). Fold enrichment was calculated as 2^−(+Ab − (−Ab))^.

### γH2A chromatin immunoprecipitation (ChIP)

We performed ChIP using rabbit antibodies that recognize phosphorylated H2A (γH2A), or no Ab control, as described above using the S9.6 antibody.

## RESULTS

### Loss of base J at telomeres leads to increased levels of TERRA

The role of base J in regulating Pol II transcription termination and the enrichment of the modified base in telomeres in *Leishmania*, suggest a role of base J in TERRA synthesis. To address this, we wanted to study the effects of J loss on TERRA synthesis in *L. major*. We have previously demonstrated the ability to titrate the levels of J in the *L. major* genome using the thymidine hydroxylase inhibitor DMOG (9,27). Increasing concentrations of DMOG led to decreasing total levels of base J, including at Pol II transcription termination sites at the end of PTUs, and defects in Pol II termination resulting in pervasive Pol II transcription of the *L. major* genome (9,27). Since it is thought that >90% of J in *Leishmania* is localized to telomeres, we first wanted to directly assess the loss of J in telomeres by DMOG. Using J-IP and telomeric qPCR, we now confirm that inhibition of J synthesis in *L. major* following growth in 5mM DMOG for 5 days, in contrast to the DMSO control, leads to the significant loss of J from telomeres as well as chromosome internal termination site represented by a convergent strand switch region (cSSR) (Figure 1A).

**Figure 1.**
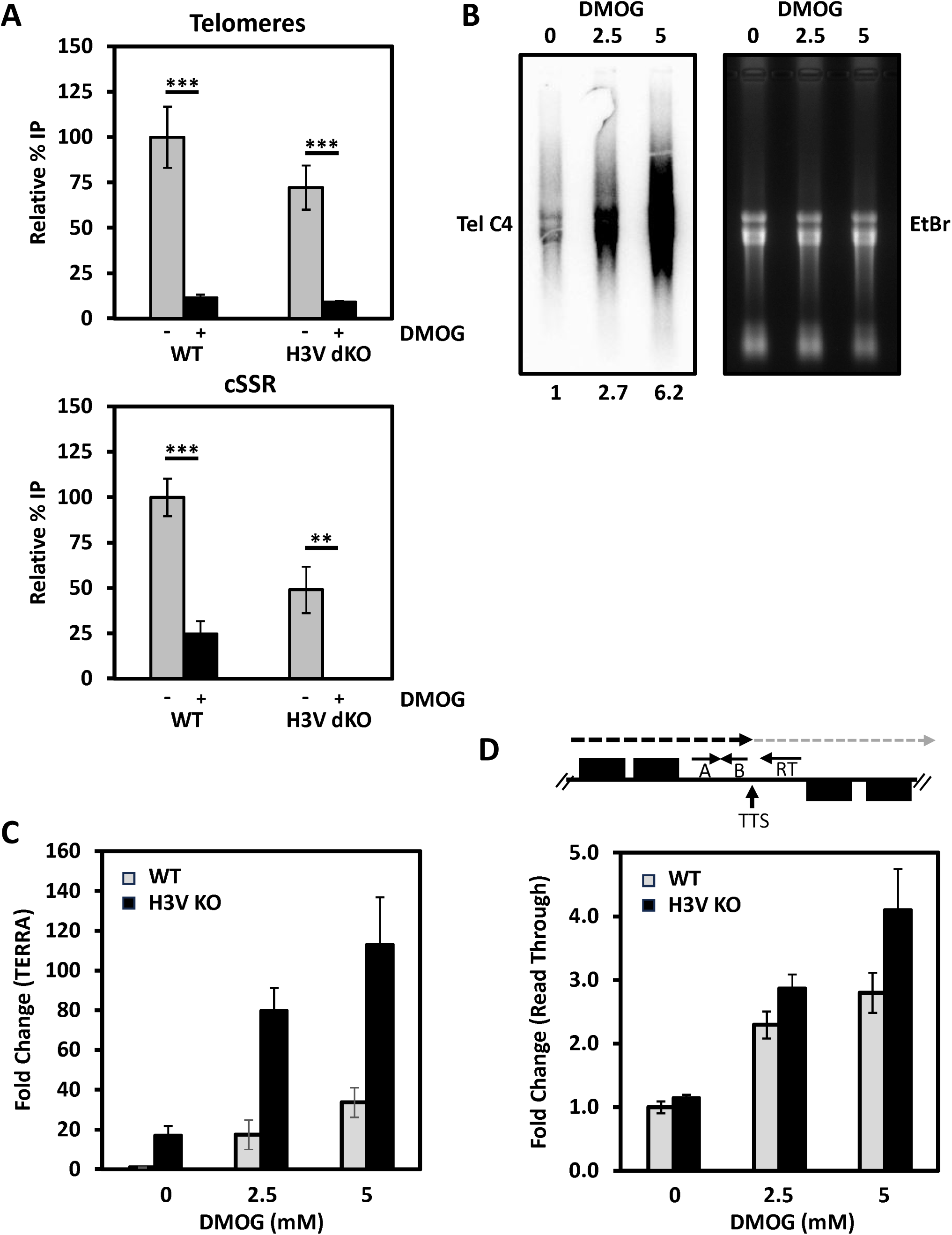
Depletion of base J results in a higher TERRA level. (A) Levels of base J in telomeres and at a chromosome internal termination site represented by convergent strand switch region (cSSR) 22.3 in *L. major* promastigotes (wild type, WT; and H3V KO) treated with and without DMOG were determined by J-IP and qPCR. The percent of DNA IP’d was calculated from three independent experiments and normalized to levels in WT set to 100. Standard deviations are shown. (B) Northern analysis of TERRA in *L. major* promastigotes treated with and without DMOG. Total RNA was isolated from cells incubated with or without DMOG and equal amounts were hybridized with the (CCCTAA)n (TelC4) probe. Right, ethidium bromide stain of the gel as a loading control. (C) Quantitation of TERRA RNA in *L. major* promastigotes (wild type, WT; and H3V KO) treated with and without DMOG by telomeric qRT-PCR. Fold changes in TERRA RNA + and – DMOG were calculated by qPCR from two biological replicates, measured in triplicate. Means and standard deviations are plotted. Fold change is relative to TERRA levels in WT cells-DMOG, normalized to tubulin qPCR. A gene specific primer against tubulin was added to the same cDNA synthesis reaction with primer RT, followed by PCR using tubulin primers. (D) Strand-specific RT-PCR analysis of readthrough transcription. Above; Schematic representation (not to scale) of cSSR 22.3 illustrating the nascent RNA species assayed by RT-PCR. The dashed arrow indicates read through transcription past the transcription termination site (TTS). Arrows indicate the location of primers utilized for strand-specific RT-PCR analysis. Below; Read through transcription on the top strand for the indicated cell lines (−/+ DMOG) was quantitated by performing site-specific cDNA synthesis using primer RT illustrated in the diagram above, followed by PCR using primers A and B. Abundance was normalized using tubulin. Fold increase in nascent readthrough RNA species relative to background levels in WT + DMSO is based on A+B qPCR analysis, normalized to tubulin qPCR. Error bars indicate the standard deviation between two biological replicates analyzed in triplicate.

To determine if the control of Pol II transcription termination by base J is involved in the transcription of telomeric repeats, we first did Northern blotting to examine the TERRA level in DMOG treated cells. Total rRNA level was detected by ethidium bromide staining as a loading control. Following treatment with 5mM DMOG, the TERRA level was significantly higher (∼ 7-fold) than the control DMSO treated cells (Figure 1B). Consistent with the effects of DMOG concentration on levels of base J (9,27), 2.5mM DMOG led to an intermediate increase in TERRA level. Analysis of the hybridization signals (including by Phosphoimager analysis) indicates that depletion of base J causes no detectable change in TERRA size (Fig 1B and Fig S1A). This can more easily be visualized in the 2.5mM DMOG treated cells (Figure 1B). To allow precise and sensitive measurement of the TERRA level, we performed telomeric specific qRT-PCR, which showed that TERRA levels increased ∼20- to ∼40-fold upon the induction of J loss in the DMOG titration (Figure 1C). We have previously demonstrated that H3V regulates J synthesis in *L. major*, but where deletion of H3V reduced J levels at termination sites at the end of PTUs with little consequence on Pol II termination (9,24). It seems that the remaining levels of J in the H3V KO were sufficient for Pol II termination at these sites. However, DMOG treatment of the H3V KO resulted in greater reduction in J levels at termination sites, even greater than DMOG treatment of wild-type parasites, that is reflected by more significant Pol II termination defects (9). First, we wanted to confirm the effects of J loss on Pol II termination at a specific termination site on chromosome 22 (cSSR 22.3). As we previously described, while the loss of J in the H3V KO does not lead to detectable levels of Pol II readthrough nascent RNA, significant levels of readthrough is seen in wild type cells treated with DMOG that is increased further in the H3V KO + DMOG (Figure 1D). However, while the loss of J in the H3V KO is not sufficient to cause Pol II termination defects at PTUs genome-wide it does lead to ∼20-fold increase in TERRA that can be increased further to >100-fold by subsequent DMOG treatment (Figure 1C). Like the effects of DMOG on base J levels and Pol II readthrough transcription at chromosome internal sites genome-wide, DMOG treatment of the H3V KO resulted in significantly higher effects on TERRA than DMOG treatment of wild type parasites. Therefore, the loss of base J in wild type *L. major* promastigotes in two different ways, DMOG inhibition of thymidine hydroxylase activity or deletion of H3V, caused an increase in levels of TERRA. The results show that base J regulates TERRA transcript levels in *L. major* promastigotes.

### Base J suppresses TERRA transcription via control of Pol II termination

Transcription of telomeres from run off Pol II transcription of telomeric PTUs in *L. major* suggest J control of TERRA is via control of Pol II termination at telomeric repeats. In *T. brucei*, TERRA is similarly transcribed via polymerase run-off during polycistronic transcription of the telomeric VSG ES PTU, but utilizing Pol I. All other systems characterized thus far utilize Pol II for TERRA synthesis. To characterize the transcription of TERRA in *L. major*, we utilized isolated nuclei to label and quantify newly synthesized RNA. As a non-radioactive label, we used 5-Ethynyl Uridine (5-EU). After the labeling reaction, we prepared total RNA from permeabilized cells, performed click chemistry with biotin, precipitated biotinylated RNA with paramagnetic streptavidin beads, reversed transcribed TERRA using the CCCTAA repeat containing TELC20 oligo (7) and quantified TERRA by qPCR (64). In control reactions, we preincubated permeabilized cells for 15 min with 1 mg/ml α-amanitin, a potent inhibitor of Pol II. The inhibition of newly synthesized TERRA, versus the Pol I transcribed Blasticidin control gene, by α-amanitin (Figure S1B) supports transcription of TERRA by Pol II in *L. major*, like yeast and human TERRA.

To explore how base J affects the TERRA level, we measured the half-life of TERRA. WT cells were treated with Actinomycin D for various lengths of time, and TERRA measured by RT-qPCR. We found that, like mammalian cells, TERRA is extremely stable. We see little degradation of TERRA at our final time point (4h)(Figure S1C). Mammalian TERRA transcripts exhibit an 3h or 8h half-life depending on their polyadenylation status, with polyadenylated TERRA having the longer half-life (71,72). It has been shown that at least a subset of the total TERRA is polyadenylated in Leishmania (48). In contrast, Tubulin is degraded with an estimated half-life like that previously measured in *L. major* promastigotes. We repeated the same experiment in Leishmania cells treated with DMOG. While we were unable to determine the actual half-life of TERRA in Leishmania, we see no evidence of increased stability of TERRA in these DMOG treated cells compared with WT cells in the four-hour time frame. If anything, there may be a slight increase in TERRA decay rate in the DMOG treated cells. Strongly suggesting that the large increase in TERRA following the loss of J is not explained by any increase in TERRA stability.

Detection of nascent telomeric RNA by RT-PCR in *L. major* extending from the final ORF of a PTU to TERRA (48), supports that at least some fraction of the total TERRA in wild-type *L. major* promastigotes is due to Pol II run-off transcription at the end of the PTU into the telomeric repeats that is controlled by base J (Figure 2A). This activity would obviously depend on the orientation of the telomeric transcription unit. Of the 72 telomeric ends in the *L. major* genome, 56 have the telomeric PTU orientated in the direction of the telomere such that readthrough transcription allows transcription of the 3’ C-rich strand of the telomere and TERRA synthesis (3’ telomere)(Fig 2B and S2). The remaining 16 chromosome ends have PTU transcription of the 5’ telomeric strand such that a distinct promoter would presumably be required for TERRA synthesis (5’ telomere). To get a better picture of TERRA synthesis across the *L. major* genome and determine if Pol II transcriptional readthrough of subtelomeric PTUs can explain the regulation of TERRA by base J, we performed telomere strand-specific RT-PCR for multiple telomeric ends representing 3’ and 5’ telomeres (Figure 2B-D). After using the CCCTAA repeat containing TELC20 oligo as a primer in the reverse direction, we used PCR primers specific for the subtelomeric region to characterize the origin of TERRA (Figure 2B). Semiquantitative RT-PCR (Figure 2B) and qRT-PCR (Figure 2C) indicate multiple chromosome ends transcribe TERRA and respond to the loss of base J using DMOG or deletion of H3V, but primarily 3’ telomeres with Pol II transcription orientation towards the telomere. Semiquantitative RT-PCR detects TERRA, in an RT dependent manner, from 3’ telomeres in WT promastigotes that is increased upon DMOG treatment and in the H3V KO (Figure 2B). In contrast, a low TERRA signal is only detected from 5’ telomeres upon DMOG treatment of WT cells and the H3V KO. Tubulin RT-PCR provides a positive control. qRT-PCR analysis of these telomeric ends, and additional 3’ telomeres, confirms the 3’ telomere specific TERRA signals and effect of DMOG on levels of TERRA (Figure 2C). If we perform RT-qPCR using primer B in the RT and PCR reactions we no longer detect any change at 3’ telomeres upon DMOG treatment in WT cells (Figure 2D), suggesting the effects previously measured at 3’ telomeres are due to changes in nascent RNA extension downstream of primer B. Thus, the effect of base J on TERRA transcription at 3’ telomeres is limited to extension of RNA into telomeric repeats. The apparent similar responses of 5’ telomeres to DMOG using primer B or telomeric repeat C20 oligo in the RT reaction suggests the changes in RNA production occurred further upstream of the subtelomeric oligo B.

**Figure 2.**
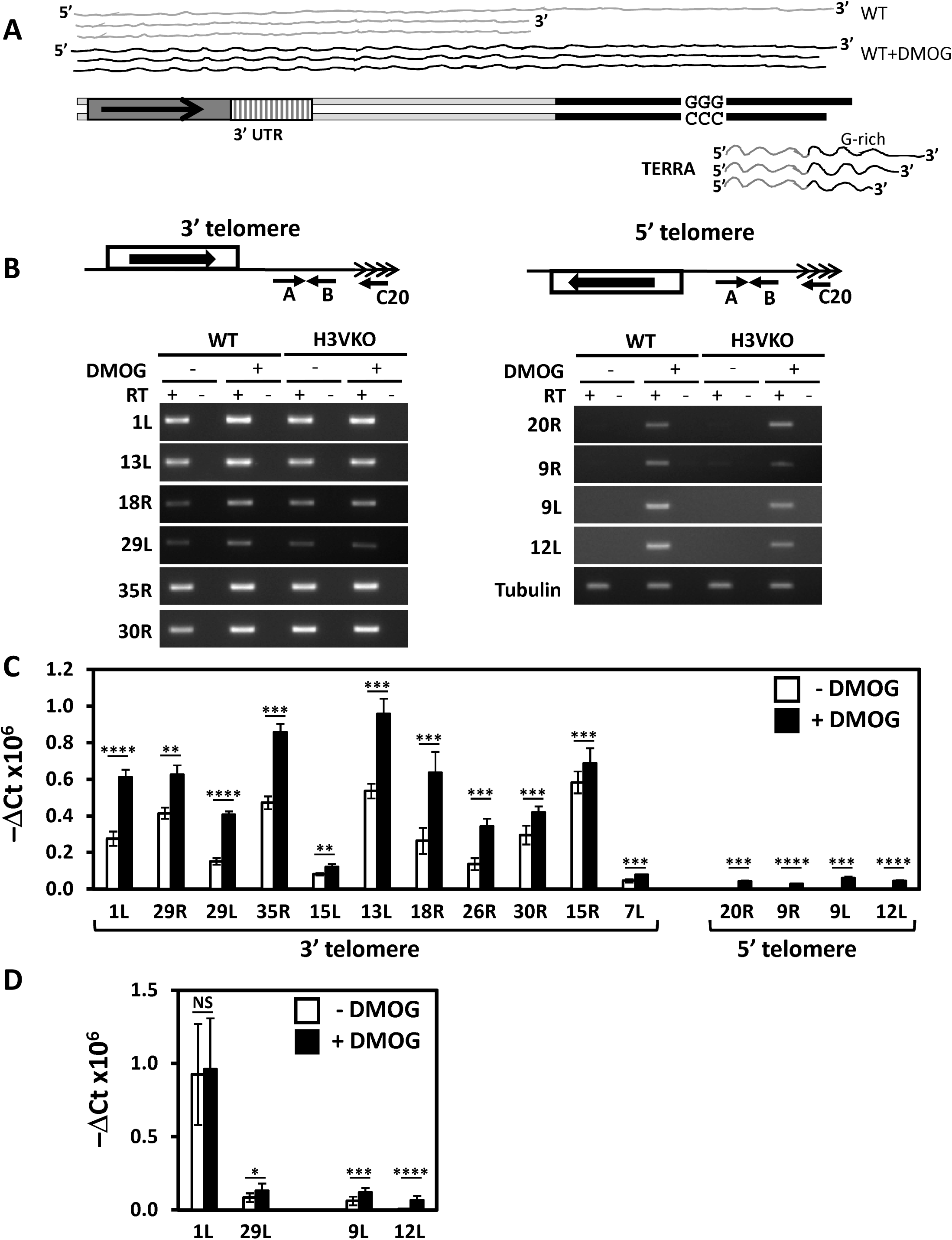
TERRA is transcribed as a Pol II readthrough transcript at multiple telomeric ends in WT and J depleted cells. (A) Schematic representation of *L. major* chromosome end and telomeric transcriptome. Telomeric repeats are in black and subtelomeric sequences in white. The final gene in the telomeric PTU is represented by the grey box and the 3’UTR indicated by the hatched box. The dashed arrow indicates the gene orientation and direction of transcription. Grey line above represents Pol II transcription of the telomere in WT cells. The black line represents the proposed extent of Pol II transcription of the subtelomeric loci upon depletion of base J. The large precursor RNA is processed to generate TERRA. The sketch is not in scale. (B) Semi-quantitative RT-PCR to detect TERRA derived from different *L. major* telomeric ends. Above, the diagram indicates the strategy used: the reverse transcription step (RT) was performed with the telomeric primer C20; the cDNA generated was used in a PCR reaction performed with oligonucleotides (primers A and B) specific for the subtelomeric region; 3’ telomeres represent regions where the final ORF is orientated towards the telomere and 5’ telomeres are where the final gene is orientated toward the internal region of the chromosome (black arrow indicated the direction of transcription). Below, shows EtBr-staining of the PCR products after fractionation in agarose gels from *L. major* promastigotes (wild type, WT; and H3V KO) treated with and without DMOG; the cDNAs used as template for the PCR were generated in the presence (+) or absence (−) of Reverse Transcriptase. The chromosome number and localization of the analyzed telomeric end on the right (R) or left (L) arm is indicated. (C) RT-qPCR analysis of TERRA expression. Ct values of TERRA levels in RT-qPCR experiments are shown for the indicated chromosome ends in WT cells treated and untreated with DMOG. TERRA Ct values were obtained after RT using TERRA C20 primer and normalized over Tubulin Ct values (2^−Δct^) obtained after RT using a Tubulin specific RT primer. Data shown represents mean +/− SD (n=3). Statistical analyses were calculated using Student’s *t*-test (∗p < 0.05, ∗∗p < 0.01, ∗∗∗p < 0.001, ∗∗∗∗p < 0.0001, ns: not significant). (D) RT-qPCR analysis of the indicated telomeric ends as in (C) but using primer B in the RT and PCR reactions.

To directly address the presence of nascent TERRA readthrough transcription product at a 3’ telomere, after using the telomere oligo in the reverse direction, we used PCR primers specific for the subtelomeric region of the right arm of chromosome 1 to amplify the nascent RNA representing Pol II run-off from the PTU into the telomeric repeats (Fig S3). While the nascent RNA is not detected in the primary PCR reaction using Tel20 and a 5’ primer specific for the 3’-UTR of the final gene of the telomeric PTU, subsequent nested PCR revealed the TERRA precursor RNA in WT cells only upon the loss of base J (+DMOG) (Fig S3). Interestingly, the levels of TERRA precursor decrease slightly with increasing levels of DMOG. These results further support that at least some fraction of the TERRA transcripts produced following the loss of base J is due to Pol II termination defects and increased readthrough RNA extending from telomeric proximal PTUs into the telomeric repeat. Taken together, the results show that base J regulation of TERRA transcript levels in *L. major* promastigotes involves the control of Pol II transcription termination from proximal PTUs.

### Base J suppression of TERRA maintains telomere integrity

Since accumulation of TERRA and associated telomeric RNA-DNA hybrids (R loops) are associate with telomere dysfunction in eukaryotes including *T. brucei* (*44–46,73,74*), we investigated the presence of R-loops and DNA damage at telomeres upon J depletion in *L. major*. TERRA is an ideal candidate to form stable RNA-DNA hybrids at telomere due the G-rich nature of its sequence composition. In *T. brucei* and *L. major* R loops are predominantly enriched in intergenic sequences and untranslated regions (70,75). Mainly excluded from Pol II-transcribed coding sequence. While little to no R loops were detected in telomeres of *L. major* promastigotes (70), we wanted to see if the loss of base J leads to detectable telomeric R loops and associated DNA damage. To determine whether the increased TERRA is associated with increased RNA-DNA hybrids at telomeres depleted of base J in *L. major*, we performed chromatin immunoprecipitation (ChIP) using an antibody (S9.6) that recognizes the RNA-DNA structure (76,77). Intergenic regions of chromosome 36 which have been previously reported to form RNA-DNA hybrids served as positive controls for the ChIP (70). We observed that wild-type cells contained the expected differing levels of RNA-DNA hybrids at the two positive control regions tested as compared to mock-treated ChIPs in which the control IgG antibody was used (Fig 3A). Also as previously described (70), we find extremely low levels of telomeric hybrids in wild-type *L. major* promastigotes, only slightly above the levels in another negative region within an ORF on chromosome 7 (ORF 07.1245). Upon DMOG treatment we see a small increase in telomeric R loops (Fig 3A). When normalized to the negative antibody control, this corresponds to a statistically significant ∼4-fold increase of telomeric R loops (Fig 3B). Interestingly, DMOG treatment leads to a slight decrease in R loops at all chromosomal internal R loop positive regions tested. To confirm that the signals detected by the antibody represent bona fide sites of RNA-DNA hybrids, ChIP samples were treated on-beads with recombinant RNase H in vitro followed by recovery and analysis of the bound DNA. However, for some unknown reason repeated RNase treatment, using enzyme from different sources, failed to have any significant effect on the signals for any locus tested (data not shown). Even though the control RNase H treatment only slightly decreased the ChIP signals, our analysis of wild-type cells accurately reflects the recent genome-wide R-loop ChIP study of *L. major* (70). Therefore, we believe our results strongly suggest a specific increase in telomeric R-loops upon the loss of base J and accumulation of TERRA.

**Figure 3.**
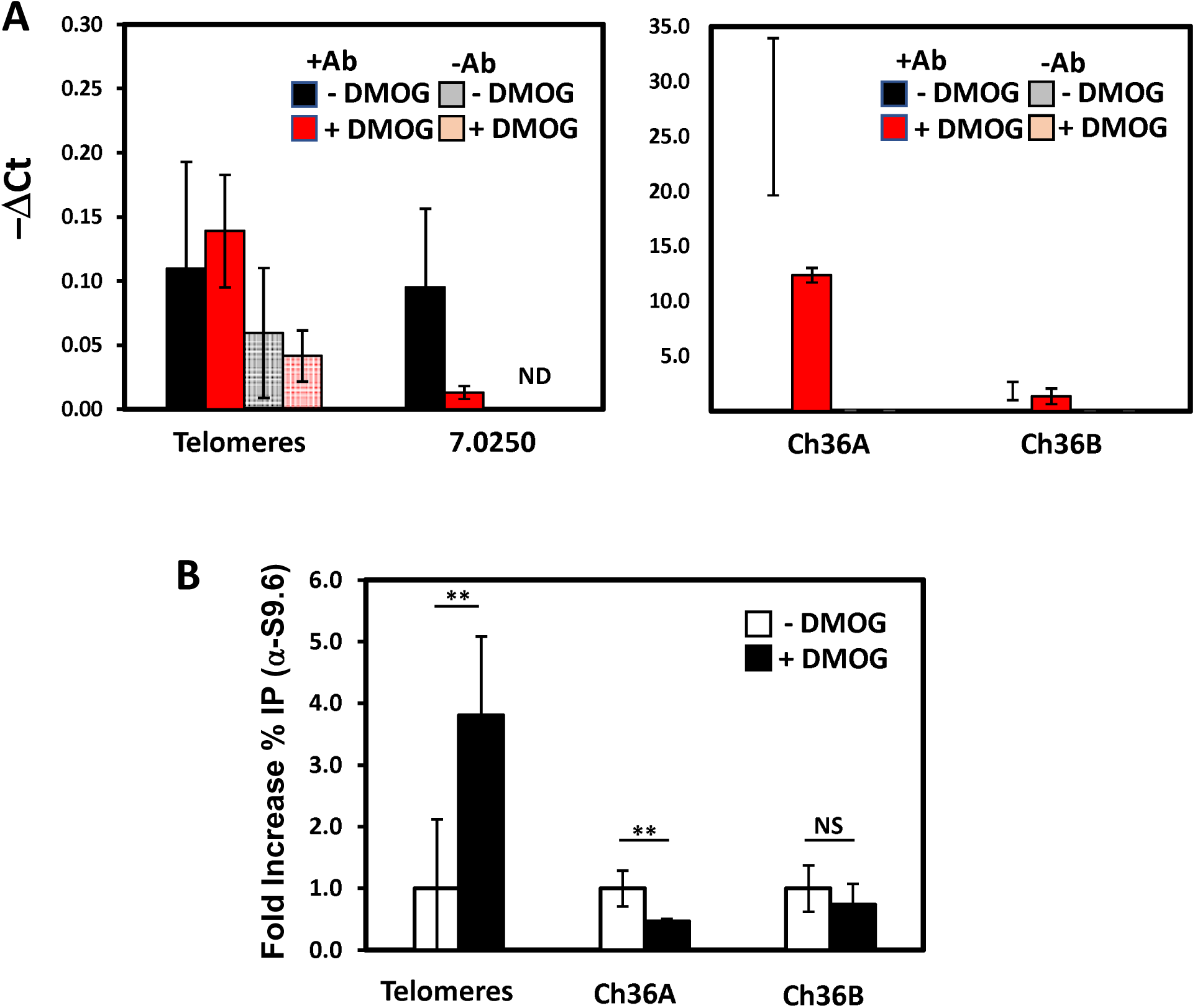
Changes in telomeric R-loops following the loss of base J. A. Detection of R-loops in L. major by immunoprecipitation with S9.6 antibody and qPCR. Immunoprecipitated DNA from RNA-DNA hybrids were amplified from telomeres and indicated genome internal regions of chromosome 7 and 36. 7.025 represents the LmjF.07.0250 ORF as R-loop negative control and Ch 36 A and B represent R-loop positive regions (70). Ct values of R-loops in each analyzed sample from cells treated with and without DMOG and IP’ed in the presence and absence of S9.6 antibody, relative to input, was plotted on each graph. Notice the different Y axis scales in the graph on the right and left. B. Fold change in the percentage of immunoprecipitated RNA-DNA hybrids in the indicated samples relative to input and normalized to the minus Ab control was plotted. Statistical analyses were calculated using Student’s *t* test (^∗^p < 0.05, ^∗∗^p < 0.01, ^∗∗∗^p < 0.001, ^∗∗∗∗^p < 0.0001, ns: not significant).

Given these effects, we surmised that TERRA amplification in *L. major*, like model eukaryotes, would cause telomere dysfunction. After treating with DMOG, we looked for formation of telomere-induced DNA damage foci using γ-H2A as a marker, which typically occur after induction of the DNA damage response at dysfunctional telomeres. The phosphorylation of H2A histones is one of the earliest responses to dsDNA breaks, helping facilitate the access of DNA-repair enzymes to nucleosomes, and γ-H2A has been used as a sensitive biomarker of DNA damage and repair in eukaryotes, including trypanosomatids (78–80). First, Western blot assays were done using a specific anti γ-TbH2A serum (44). As the positive control to test if LmH2A is indeed phosphorylated in response to DNA damage, we exposed WT promastogotes to phleomycin, a DNA-damage agent that induces DNA double-strand breaks (81–83). Exponentially growing control promastigotes were incubated with phleomycin, and aliquots were harvested after 0, 1, 2, and 6 hr. The time-course experiment shows a clear increase in the intensity of the 15-kDa band, which corresponds to yH2A protein, in response to increase DNA breaks induced by phleomycin (Figure 4A). These data suggest that LmH2A is phosphorylated in response to DNA damage and further confirm that the anti-γ-TbH2A antibody is specific to the phosphorylated form of LmH2A.

**Figure 4.**
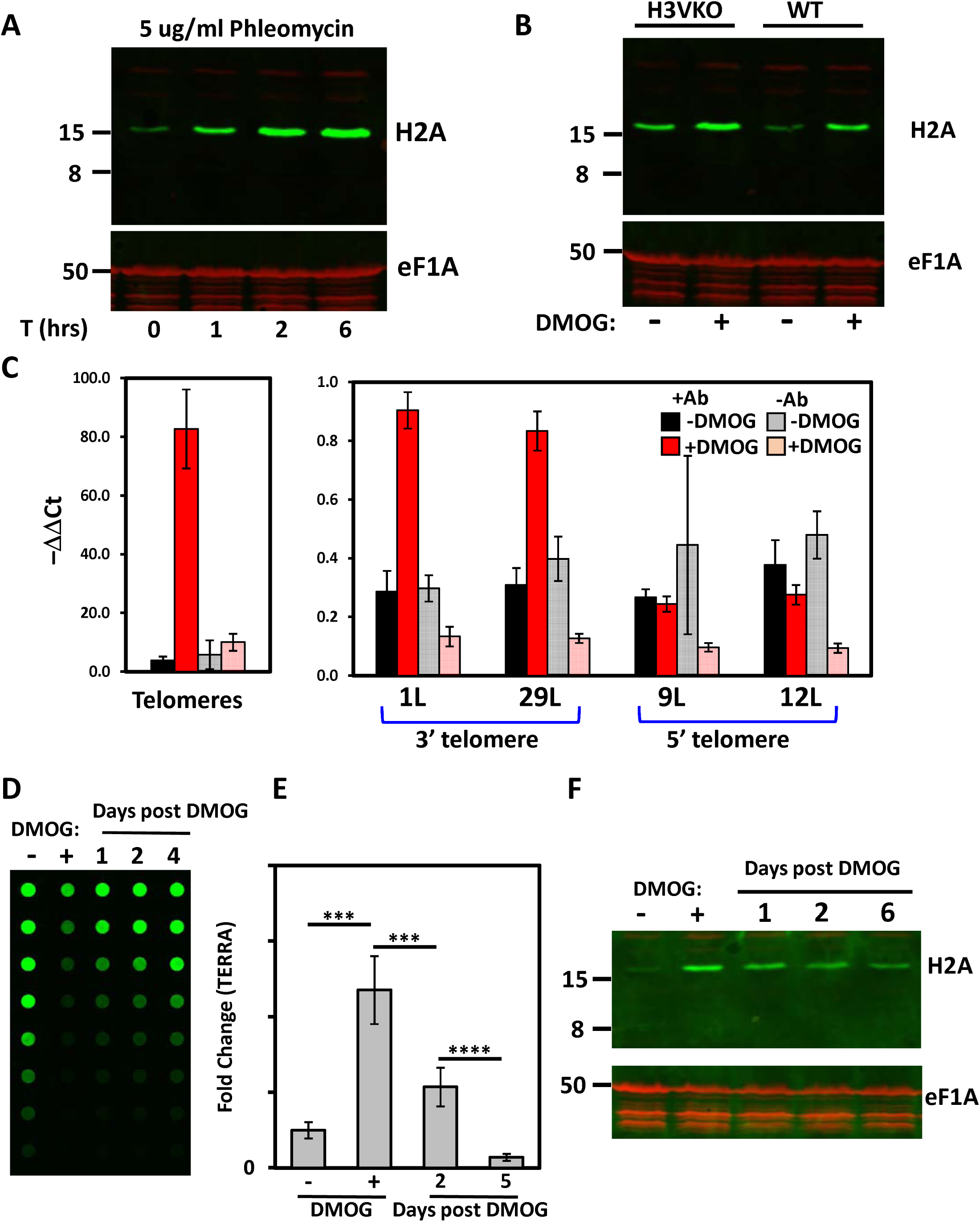
Loss of base J in *L. major* results in increased double strand DNA breaks at 3’ telomeres. A. DNA damage in *L. major* causes histone H2A phosphorylation. WT *L. major* promastigote cells were incubated with 5ug/ml of phleomycin for the indicated periods of time and extracts were analyzed by western blot with anti-γH2A antibody. EF1a was used as a loading control. B. Loss of base J leads to increased phosphorylation of histone H2A. Extracts from WT and H3V KO cells treated with and without DMOG were analyzed by western blot with anti-γH2A antibody. EF1a was used as a loading control. Western blot is representative of 3 independent experiments. C. ChIP experiments were performed using the antibody recognizing γH2A or without Ab (as a negative control). DNA isolated from the ChIP products were analyzed by qPCR to determine the levels of DSBs at total telomeres and the indicated specific telomeric regions representing 3’ and 5’ telomeres as in Figure 2C. Enrichment (ChIP/input) was calculated from three independent experiments. Standard deviations are shown as error bars. (D-F.) The TERRA and DNA damage response induction is dynamic. WT cells were incubated 6 days with DMOG, then washed to eliminate residual DMOG and fresh media was added to harvest cells at the indicated times. D. J dot blot analyzing the level of base J in genomic DNA. E. Quantitation of TERRA RNA levels determined in the indicated samples by telomeric qRT-PCR as in Figure 1C. Fold changes in TERRA RNA were calculated by qPCR from two biological replicates, measured in triplicate. Standard deviations are shown as error bars. P values of unpaired t-tests were shown for several pairs of values. F. Extracts from the indicated samples were analyzed by western blot with anti-γH2A antibody. EF1a was used as a loading control. Western blot is representative of 3 independent experiments.

The ability of *L. major* promastigotes to phosphorylate histone H2A in response to loss of base J was assessed by western blot. Using γ-TbH2A antibody on proteins extracted from WT parasites grown with and without DMOG showed a corresponding elevation in the levels of γ-H2A following DMOG treatment (Figure 4B). While H3V KO cells have similar levels as WT+DMOG, DMOG treatment of the H3VKO leads to a further increase in the levels of γ-H2A. This finding suggests that increased TERRA and corresponding telomeric R-loops in the base J mutants is associated with DNA damage accumulation. To explore specific DNA damage response in telomeres we performed a Chip-PCR analysis using the γ-H2A antibody to compare levels of DNA damage in WT cells treated with DMOG at different telomeric ends. More telomeric DNA was seen to associate with γ-H2A upon DMOG treatment (Figure 4C), indicating that the loss of J indeed resulted in an increased amount of DNA damage at the telomere. We also see increased DNA damage at 3’ telomeres compared to 5’ telomere (Figure 4C), consistent with the specific increases in TERRA expression (Figure 2B and C). These observations suggest more DNA damage at the telomere upon the loss of base J and corresponding increased TERRA transcription. Therefore, base J has a role in maintaining telomere integrity presumably via control of TERRA synthesis and R loop formation.

### TERRA and telomere damage control by base J is dynamic

To gain further insight regarding TERRA modulation by base J in response to DMOG, we evaluated whether TERRA induction, and associated DNA damage response, was a dynamic event. We have previously demonstrated in *T. brucei* that following the inhibition of base J syntheses removal of the DMOG pressure leads to rescue of J levels in the genome and reversal of Pol II transcriptional regulatory phenotypes in 4-5 days. To explore the dynamic nature of J function in *L. major*, we treated cells with DMOG for 5 days, eliminated the residual DMOG, added fresh medium and harvested cells after 1, 2, 4 and 6 days had elapsed. Like the response in *T. brucei*, we see recovery of J levels after removal of DMOG in *L. major* promastigotes, with ∼50% WT levels of J restored by day 4 (Figure 4D). Importantly, the re-synthesis of base J following DMOG removal restored TERRA levels by day 5 (Figure 4E), confirming that the impaired level of TERRA is J dependent and that TERRA induction was a dynamic event. We found that LmH2A phosphorylation is also dynamic and that over time, once base J levels are rescued and TERRA levels recover, levels of phosphorylated H2A reduce to approximately WT levels by day 6 (Figure 4F). Presumably this represents reversal of DNA damage at 3’ telomeres. Interestingly, the growth defects of parasites grown in the presence of 5mM DMOG also appear to be reversed following removal of DMOG and recovery of base J synthesis. The dynamic nature of base J synthesis and phenotypes measured supports direct involvement of base J in TERRA transcription and telomere dysfunction.

### Control of metacyclogenesis by base J regulated Pol II transcription

As mentioned above, treatment of wild type *L. major* with DMOG decreases J levels below a certain cutoff leading to readthrough transcription, and the degree of readthrough transcription in these cells is stimulated by further loss of J upon the deletion of H3V (9)(Fig 1D). We further show here that altering the levels of J at the end of telomeric PTUs using DMOG and the H3V mutant lead to similar corresponding changes in levels of TERRA from readthrough transcription and telomeric DNA damage in *L. major* promastigotes. Observation of these cells under the microscope suggested the loss of J and Pol II readthrough leads to morphological changes associated with the formation of non-dividing metacyclics. To address this further, we quantitated the levels of metacyclics in cultures of *L. major* promastigotes with varying levels of base J by negative selection with peanut agglutinin. Initially, we studied differences in metacylics generated during growth of WT and H3V KO parasites to stationary phase. As shown in Fig 5A, we detect an increase in numbers of metacyclic promastigotes (PNA-) in the H3V KO compared to WT parasites. Next, we quantitated the levels of metacyclics generated during log phase growth of *L. major* parasites treated with DMOG. As expected, the numbers of metacyclic parasites in log phase are very low, below 1%, in wild-type cultures (Fig 5A). However, we detect an increase in numbers of metacyclic promastigotes upon the loss of J in wild-type cells treated with DMOG, that increases slightly further upon the additional loss of J in H3V+DMOG (Figure 5B), a phenomenon that closely mimics the increases in Pol II readthrough transcription at the end of PTUs (9)(Fig 1D), and corresponding increased levels of TERRA (Figure 2B and C) and DNA damage (Fig 4B). The results suggest that the control of Pol II termination by base J, and expression of TERRA and control of telomere dysfunction is associated with the control of the differentiation of *L. major* promastigotes to the non-dividing infective metacyclic life stage.

**Figure 5.**
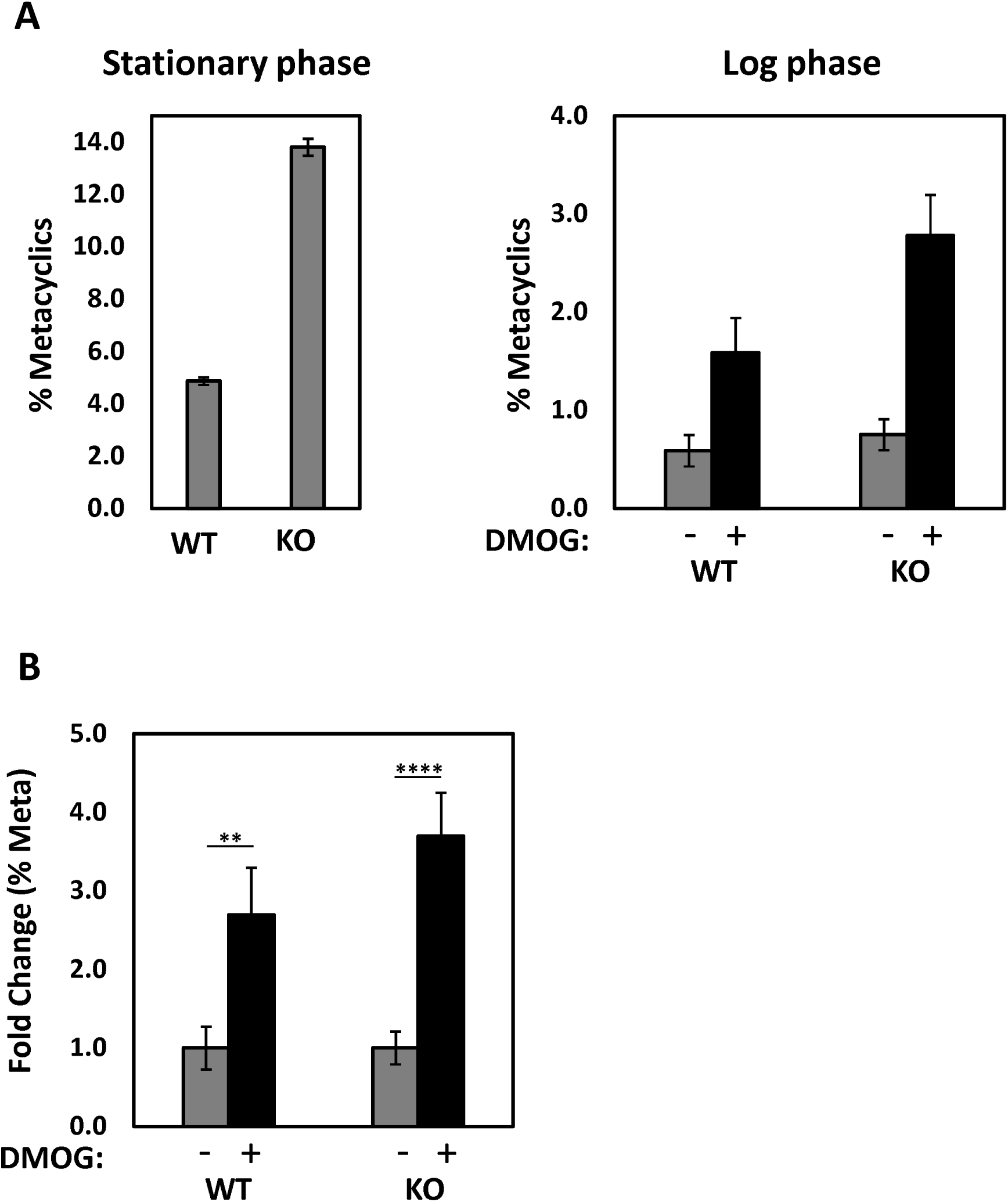
Base J regulation of metacyclogenesis. Metacyclic promastigotes were enriched by negative selection with peanut agglutinin of WT and H3V KO procyclic promastigotes in (A) stationary phase without DMOG treatment and (B) Log phase following treatment without and with 5mM DMOG for 5 days. (C) Fold change in percent of cells remaining are expressed relative to the WT minus DMOG control (set to one). WT, wild-type; KO, H3V KO. Means and standard deviations from at least three independent experiments are plotted.

## DISCUSSION

In this study, we establish a direct mechanistic link between the hypermodified DNA base J, Pol II transcription termination, telomeric RNA (TERRA) synthesis, telomere integrity, and developmental progression in *Leishmania major*. While base J has previously been characterized as an epigenetic mark required for proper termination at polycistronic transcription units (PTUs), the present study demonstrates that this regulatory system extends to chromosome ends, where base J suppresses inappropriate TERRA synthesis and preserves telomere stability. It is becoming clear that TERRA expression must be tightly regulated in trypanosomatid parasites; where induction of TERRA expression can fuel the DNA damage response at chromosome ends during telomere dysfunction. These results provide a coherent framework that helps explain why the JBP1 KO was unable to be generated and the essential nature of base J in *Leishmania*.

A central finding of this work is that TERRA in *L. major* is synthesized by Pol II and arises primarily from transcriptional readthrough of subtelomeric PTUs oriented toward chromosome ends. This mechanism resembles Pol II–dependent TERRA synthesis in yeast and mammalian systems, but contrasts with *Trypanosoma brucei*, where TERRA is generated by Pol I run-off transcription from the active VSG expression site. Thus, kinetoplastids employ distinct polymerase contexts for telomeric transcription, and in *L. major*, TERRA production is tightly coupled to Pol II termination fidelity. Our data demonstrate that depletion of base J, either through inhibition of thymidine hydroxylases with DMOG or in the absence of H3V, results in substantial upregulation of TERRA from multiple telomeres, particularly those with PTUs transcribed toward the telomeric repeats. Importantly, increased TERRA is not explained by enhanced transcript stability, indicating that base J acts primarily at the level of transcription.

These results reinforce the model that base J functions as a critical determinant of Pol II termination at PTU boundaries, including those proximal to telomeres. Given that >90% of base J localizes to telomeric regions in *Leishmania*, our findings suggest that one major biological role of this DNA modification is to prevent pervasive transcription into repetitive telomeric DNA. Telomeres represent structurally and functionally specialized chromosomal domains; uncontrolled transcription into telomeric repeats may have amplified consequences compared with internal genomic regions. The pronounced increase in TERRA following J depletion—reaching up to ∼100-fold under combined H3V loss and DMOG treatment—highlights the sensitivity of telomeric transcription to termination defects.

Studies on the telomere position effect on gene silencing in yeast has indicated subtelomeres can control transcription in many ways, primarily involving control of chromatin structure (84,85). These mechanisms may play a role in telomere transcription of TERRA. For example, Sir-complex-dependent chromatin modifications have been shown to be a key regulatory mechanism limiting the global levels of TERRA in yeast. In mammalian cells, promoters have been found in CpG-islands within subtelomeric regions of human chromosomes and where TERRA transcription initiation is controlled by the chromatin state and DNA methylation (52,86,87). Subtelomeres in yeast can also help limit TERRA transcript levels by regulating Rap1-mediated pathways (88). Rap1 mutants have increased levels of TERRA, presumably by the loss of transcription repression by reduced recruitment of Sir complex components as well as by decreased Rat1-dependent degradation (88). Increased TERRA transcription, and formation of telomeric R-loops, was detected in *T. brucei* cells depleted of TbRap1 (44). TbRap1 is also essential for cell viability and VSG silencing and VSG switching (44,89,90). Furthermore, chromatin modifications play a role in silencing transcription of TERRA in the silent VSG ESs by the native telomeric Pol I promoter as well as any leaky Pol II elongation from a distant promoter. While Pol II termination defects in *T. brucei* leads to Pol II transcription into all the telomeric PTUs, TERRA synthesis is still restricted to the active VSG ES (7). Pol II readthrough is terminated just before the telomeric repeats in the silent VSG ESs presumable by the repressive telomeric chromatin. Reb1, an essential transcription factor in yeast, can bind subtelomeres and control TERRA transcription directly, where the reb1 mutant has 75-fold higher levels of TERRA (43). It is presumed that Reb1 represses TERRA levels by a roadblock transcriptional termination mechanism, independent of Sir-complex regulation of chromatin. While base J may provide some level of steric block for Pol II elongation in trypanosomatids, we believe the main mechanism involves control of Pol II-CTD phosphorylation and torpedo dissociation of the polymerase from the DNA template at termination sites (5–7). Consistent with this idea, analysis of the hybridization signals on the northern blot indicates that depletion of base J causes no detectable change in TERRA size. This suggests that the increase in TERRA is not due to effects of J on the extent of Pol II elongation within the telomeric repeat but rather help control TERRA levels by termination of transcription prior to the telomeric repeats.

Elevated TERRA levels in J-deficient parasites are accompanied by increased telomeric RNA–DNA hybrids and accumulation of γ-H2A, consistent with enhanced DNA damage at chromosome ends. In wild-type promastigotes, telomeric R-loops are minimal, but J depletion results in a reproducible and significant increase in the R-loop IP signal. The correlation between TERRA induction, telomeric R-loops, and γ-H2A enrichment at telomeres strongly supports a model in which base J suppresses telomere instability by limiting transcriptional run-off and DNA-RNA hybrid formation. In other eukaryotes, excessive TERRA and persistent R-loops promote replication stress, double-strand breaks, and recombination at telomeres, threatening genome integrity. Our data indicate that similar principles operate in *L. major*, where loss of J-mediated termination control destabilizes telomeres and activates a DNA damage response.

Notably, these phenotypes are dynamic and reversible. Restoration of base J levels following removal of DMOG leads to normalization of TERRA abundance, reduction of γ-H2A, and recovery of growth. This reversibility argues strongly for a direct role of base J in regulating telomeric transcription and DNA damage rather than an indirect or secondary effect of prolonged cellular stress. The rapid recovery of telomere-associated phenotypes further emphasizes that base J–dependent transcription termination is a continuously required process for maintaining telomere homeostasis.

An unexpected but biologically significant consequence of J depletion is enhanced differentiation of promastigotes into the infectious metacyclic stage. Increased metacyclogenesis correlates with the degree of Pol II readthrough, TERRA upregulation, and telomeric DNA damage. A link between J, TERRA and metacyclogenesis has previously been suggested by noticing a potential decrease in J levels localized to telomeres upon differentiation of promastigotes to metacyclics (37) while levels of TERRA increase (37,48). Together, these observations suggest that disruption of epigenetic termination control and telomere stability influences developmental progression. Because transcription in trypanosomatids is largely constitutive and polycistronic, regulation at the level of transcription termination provides a mechanism for modulating gene expression programs without conventional promoter-based control. It is plausible that telomere dysfunction or pervasive transcription acts as a stress signal that promotes differentiation, or that altered termination at specific subtelomeric loci directly derepresses genes involved in developmental transitions. In either case, our data position base J–mediated termination as a key link between genome stability and life cycle regulation.

Taken together, these findings support a model in which base J, through recruitment of termination machinery and enforcement of Pol II boundary control, prevents transcriptional encroachment into telomeric repeats. In the absence of base J, Pol II readthrough increases TERRA synthesis, promotes R-loop accumulation, induces telomeric DNA damage, decreases cell viability, and biases parasites toward metacyclogenesis. Thus, the essentiality of base J in *Leishmania* likely reflects the combined consequences of genome-wide transcriptional dysregulation and telomere-specific instability.

More broadly, this work highlights an intimate connection between epigenetic DNA modification, transcription termination, noncoding RNA production, chromosome-end stability, and developmental regulation in an early-diverging eukaryote. Future studies should define how base J deposition is targeted and developmentally regulated at telomeres, determine whether TERRA directly drives differentiation or primarily mediates telomere dysfunction, and dissect the molecular pathways linking transcriptional readthrough to DNA damage signaling. Understanding how base J coordinates transcription boundaries with telomere biology may reveal conserved principles of genome regulation and provide new insight into parasite adaptation during the life cycle of *Leishmania major*. Finally, the ability of DMOG to alter TERRA levels in vivo without the need of other functional genetic approaches, provides a useful tool to elucidate TERRA function and synthesis in Leishmania. Regulating J synthesis may provide a unique way to manipulate TERRA specifically in vivo without simultaneously altering additional telomere components. Providing a unique resource not available in other eukaryotic systems.

## Funding

This work was supported by the National Institutes of Health (grant number R01AI109108) to R.S. The content is solely the responsibility of the authors and does not necessarily represent the official views of the National Institutes of Health.

## Conflict of interest

The authors declare that they have no conflicts of interest with the contents of this article.

## Acknowledgment

We would like to thank David Reynolds for the initial experiments linking J loss with stimulating metacyclogenesis. We also thank Bibo Li for providing the anti-γH2A antisera.

## Notes

### Competing Interest Statement

The authors have declared no competing interest.

### Summary of Updates

This updated version contains additional and new data concerning TERRA formation, R-loops, DNA damage and metacyclogenesis in L major

