## Supplemental Figures for "Epigenetic regulation of TERRA transcription, R-loop formation, telomere integrity and metacyclogenesis by base J in *Leishmania major*"

### Slide 1
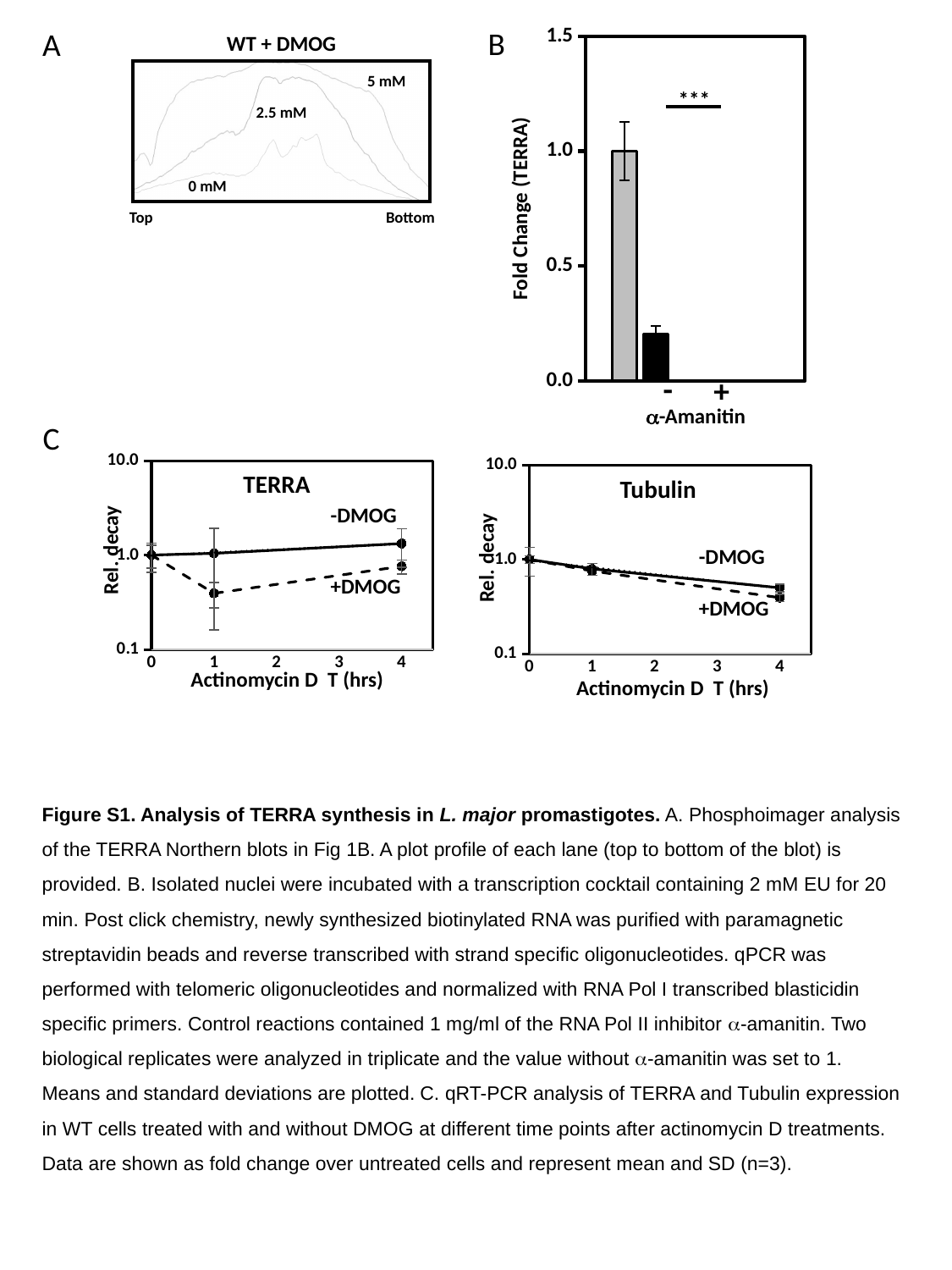

B
A
[unsupported chart]
WT + DMOG
5 mM
***
2.5 mM
0 mM
Fold Change (TERRA)
Top
Bottom
-
+
α-Amanitin
C
#### Chart
| Category |
|---|
#### Chart
| Category | | |
|---|---|---|TERRA
Tubulin
-DMOG
Rel. decay
-DMOG
Rel. decay
+DMOG
+DMOG
Actinomycin D T (hrs)
Actinomycin D T (hrs)
Figure S1. Analysis of TERRA synthesis in L. major promastigotes. A. Phosphoimager analysis of the TERRA Northern blots in Fig 1B. A plot profile of each lane (top to bottom of the blot) is provided. B. Isolated nuclei were incubated with a transcription cocktail containing 2 mM EU for 20 min. Post click chemistry, newly synthesized biotinylated RNA was purified with paramagnetic streptavidin beads and reverse transcribed with strand specific oligonucleotides. qPCR was performed with telomeric oligonucleotides and normalized with RNA Pol I transcribed blasticidin specific primers. Control reactions contained 1 mg/ml of the RNA Pol II inhibitor α-amanitin. Two biological replicates were analyzed in triplicate and the value without α-amanitin was set to 1. Means and standard deviations are plotted. C. qRT-PCR analysis of TERRA and Tubulin expression in WT cells treated with and without DMOG at different time points after actinomycin D treatments. Data are shown as fold change over untreated cells and represent mean and SD (n=3).

### Slide 2
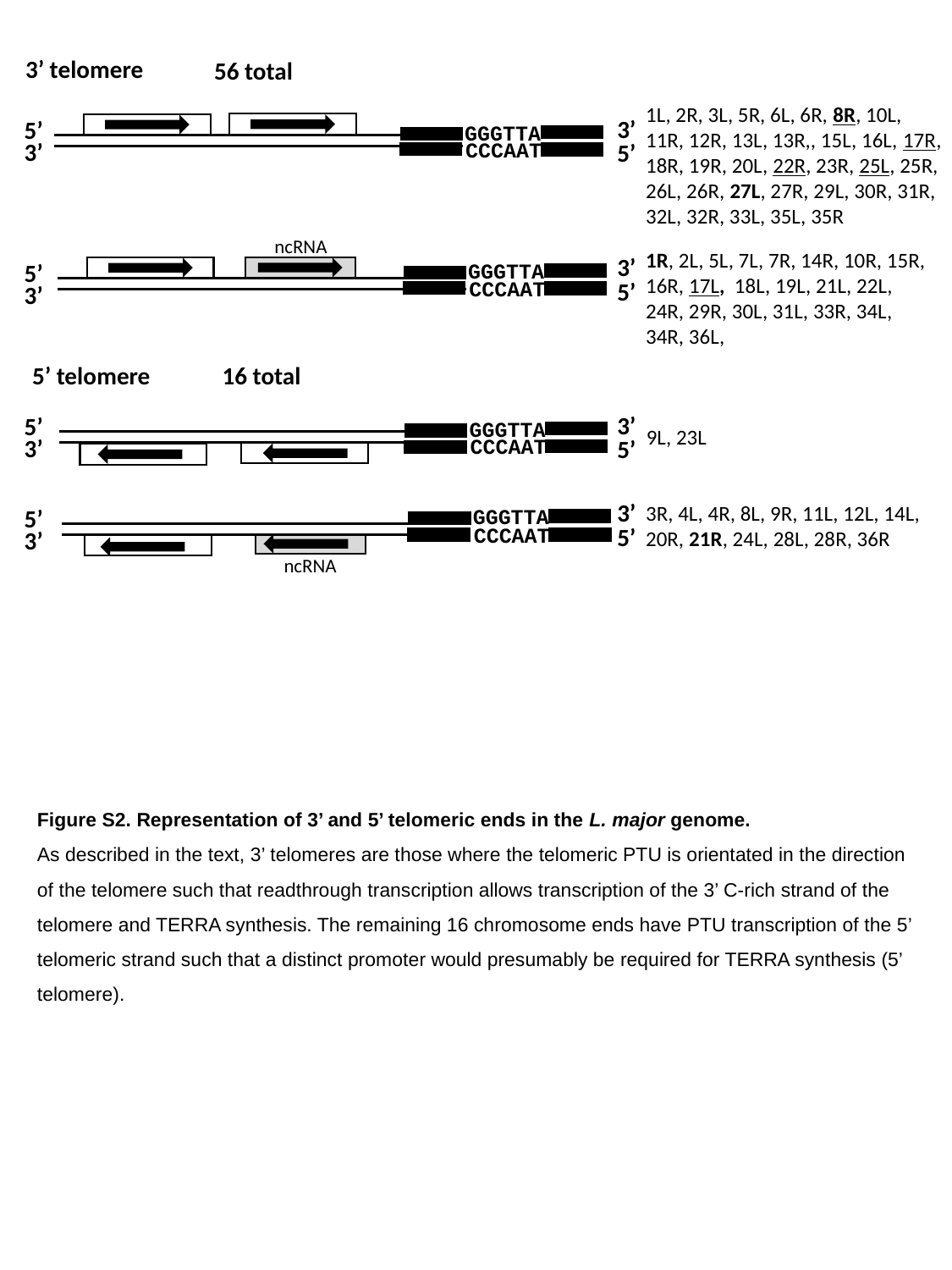

3’ telomere
56 total
1L, 2R, 3L, 5R, 6L, 6R, 8R, 10L, 11R, 12R, 13L, 13R,, 15L, 16L, 17R, 18R, 19R, 20L, 22R, 23R, 25L, 25R, 26L, 26R, 27L, 27R, 29L, 30R, 31R, 32L, 32R, 33L, 35L, 35R
3’
5’
GGGTTA
3’
CCCAAT
5’
ncRNA
1R, 2L, 5L, 7L, 7R, 14R, 10R, 15R, 16R, 17L, 18L, 19L, 21L, 22L, 24R, 29R, 30L, 31L, 33R, 34L, 34R, 36L,
3’
5’
GGGTTA
CCCAAT
5’
3’
5’ telomere
16 total
3’
5’
GGGTTA
9L, 23L
3’
CCCAAT
5’
3’
3R, 4L, 4R, 8L, 9R, 11L, 12L, 14L, 20R, 21R, 24L, 28L, 28R, 36R
5’
GGGTTA
5’
CCCAAT
3’
ncRNA
Figure S2. Representation of 3’ and 5’ telomeric ends in the L. major genome.
As described in the text, 3’ telomeres are those where the telomeric PTU is orientated in the direction of the telomere such that readthrough transcription allows transcription of the 3’ C-rich strand of the telomere and TERRA synthesis. The remaining 16 chromosome ends have PTU transcription of the 5’ telomeric strand such that a distinct promoter would presumably be required for TERRA synthesis (5’ telomere).

### Slide 3
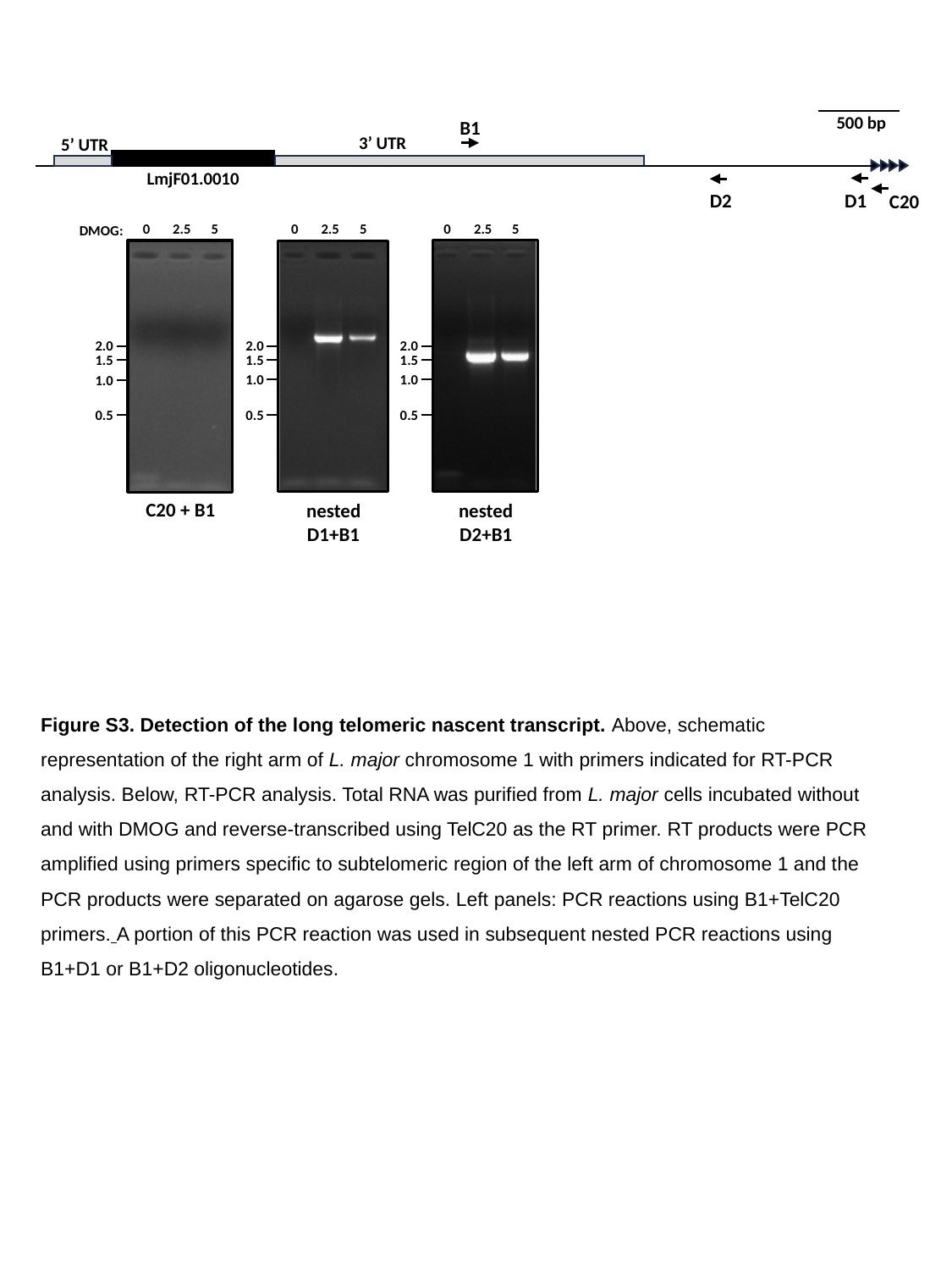

500 bp
B1
3’ UTR
5’ UTR
LmjF01.0010
D2
D1
C20
0
2.5
5
0
2.5
5
0
2.5
5
DMOG:
2.0
1.5
1.0
0.5
2.0
1.5
1.0
0.5
2.0
1.5
1.0
0.5
C20 + B1
nested
D1+B1
nested
D2+B1
Figure S3. Detection of the long telomeric nascent transcript. Above, schematic representation of the right arm of L. major chromosome 1 with primers indicated for RT-PCR analysis. Below, RT-PCR analysis. Total RNA was purified from L. major cells incubated without and with DMOG and reverse-transcribed using TelC20 as the RT primer. RT products were PCR amplified using primers specific to subtelomeric region of the left arm of chromosome 1 and the PCR products were separated on agarose gels. Left panels: PCR reactions using B1+TelC20 primers. A portion of this PCR reaction was used in subsequent nested PCR reactions using B1+D1 or B1+D2 oligonucleotides.
